# A deep learning approach for building multiple trees classification

**DOI:** 10.1101/781252

**Authors:** Nadia Tahiri

## Abstract

Each gene has its own evolutionary history which can substantially differ from the evolutionary histories of other genes. For example, some individual genes or operons can be affected by specific horizontal gene transfer or hybridization events. Thus, the evolutionary history of each gene should be represented by its own phylogenetic tree which may display different evolutionary patterns from the species tree, or Tree of Life, that represents the main patterns of vertical descent. Here, we present a new efficient method for inferring single or multiple consensus trees and supertrees for a given set of phylogenetic trees (i.e. additive trees or X-trees). The output of the traditional tree consensus methods is a unique consensus tree or supertree. Here, we show how Machine Learning (ML) models, based on some interesting properties of the Robinson and Foulds topological distance, can be used to partition a given set of trees into one (when the data are homogeneous) or multiple (when the data are heterogeneous) cluster(s) of trees. We adapt the popular *Accuracy, Precision, Sensitivity*, and *F*_1_ scores to the tree clustering. A special attention is paid to the relevant, but very challenging, problem of inferring alternative supertrees that are built from phylogenies defined on different, but mutually overlapping, sets of species. The use of an approximate objective function in clustering makes the new method faster than the existing tree clustering techniques and thus suitable for the analysis of large genomic datasets.

## Introduction

In recent years, the next-generation sequencing (NGS) has revolutionized systematic biology and molecular ecology. NGS is a fast, reliable and affordable sequencing technique which produces millions of DNA sequence reads in a single run (Mardis 2008; Glöckle et al. 2014). The evolutionary biology applies tree inferring methods to the aligned sequences in order to reconstruct the species phylogeny which represents the evolution of the species under study. The most popular tree inferring methods nowadays include Neighbor-Joining (Saitou and Nei, 1987), PhyML (Guindon et al. 2010) and RAxML (Stamatakis 2014). Most of the conventional trees inferring methods generate one candidate tree for a given set of input data. However, the topologies of gene trees representing the evolution of different genes can be substantially different due to possible horizontal gene transfer, hybridization or intragenic and intergenic recombination events they may undergo. Each gene tree depicts a unique evolutionary history, which is connected to the species tree but often considerably diverges from it (Szöllősi et al. 2014). In order to produce a reliable species phylogeny, the related gene trees should be merged, while minimizing the topological conflicts presented in them (Maddison et al. 2007). Two scenarios are envisaged here: 1) Merging trees defined on the same set of species, which are usually associated with the tree leaves (the case of consensus trees), and 2) Merging trees defined on different, but mutually overlapping, sets of species (the case of supertrees).

A large variety of methods have been proposed to resolve the problem of reconciliation of multiple trees in order to reconstruct a species tree. In case of consensus trees, the most known types of consensus trees are the strict consensus tree, the majority-rule consensus tree and the extended majority consensus tree (Bryant 2003; Felsenstein 2004). Three main methods have been proposed to synthesize collections of small phylogenetic trees with incomplete taxon overlap into comprehensive supertrees, which include all taxa found in the input trees. The most known methods are Matrix Representation with Parsimony (MRP) (Baum 1992; Ragan 1992), parsimony supermatrix (Driskell et al. 2004; Siccarelli et al. 2006) and strict supertree (Sanderson et al. 1998). The MRP method is an amply used supertree method, which analyzed and inferred separately different systematic data sets. The trees derived from these independent analyses are used to produce a single MRP matrix and then this matrix was analyzed to reconstruct the supertree of all source taxa (de Queiroz and Gatesy, 2007). Another way is a parsimony supermatrix approach, this last one concatenates all systematic characters into a single phylogenetic matrix and then by analyzing all the characters simultaneously to compute supermatrix tree. The strict supertree represents the bipartitions which agree with all bipartitions present in phylogenetic tree sources, the rest of bipartitions are indicated by multifurcations. The strict supertree is often no resolved tree.

The implementation of the famous project “Tree of Life” (ToL) intended for inference of the largest possible species phylogenetic tree became feasible because of collaborative efforts of biologists and nature enthusiasts from around the world (1). The approach adopted by the project organizers allows for a gradual reduction of the complex tree reconstruction problem into several sub-problems and then for merging the obtained results. Indeed, such an approach produces thousands of small trees which should be combined in order to assemble ToL. The problem is twofold: first, we have to infer small sub-trees of ToL (i.e., often gene trees) defined on in this project only on the same sets of taxa; second, we have to merge these small trees into one or several large consensus phylogenies using a consensus trees reconstruction algorithm (i.e., Phylip Package of (2), which allows for combining trees inferred for same, sets of taxa (3). In this context, the application of a tree or a consensus-tree clustering method would allow for providing alternative evolutionary scenarios for several sub-trees of ToL.

Moreover, biological dataset is often heterogeneous, i.e., source of the dataset can be Deoxyribonucleic acid, Ribonucleic acid or protein, which the length is variable and it is more complicated to interpret. The heterogeneous of biological dataset implies completeness and difficulty of data interpretation. Thus, it will a good testbed for Deep Learning (DL) techniques. The superiority of DL approach is acknowledged for prediction. We aim to exploit the phylogenetic relationship (via distance between trees) to enable adopting the Convolutional Neural Network (CNN) in DL architecture. The operation is based on the pairwise Robinson and Foulds (RF) topological distances (4) stored in matrix structure.

In this paper, we describe a new tree clustering method based on CNN. We illustrated the results of simulations by loss and accuracy functions. Finally, we apply this strategy to four real datasets of (5).

## Phylogeny

In this section, we give some terminology which is necessary to describe our new approach.

### Trees

Figure 1 presents an example of phylogenetic tree defined on a set of five species (i.e., Human, Orang-Outan, Mouse, Rat, and Bird). The trees are generally (in Bioinformatic) defined by their Newick strings (2) and could be unresolved or resolved. In the first case (i.e., unresolved trees), the phylogenetic tree lack of knowledge and it is representing by multifurcation, more than three branches by internal node. Figure 1 illustrates one resolved phylogenetic tree, i.e., each internal node has three branches. Figure 1 displays that Human and Orang-Outan are closed and the same concept for Rat and Mouse. Finally, we observed that the Bird is evolutionary different of these two clades ({Human, Orang-Outan} and {Rat, Mouse}). It means that Bird was considered like an outlier in this study.

11

**Fig. 1.**
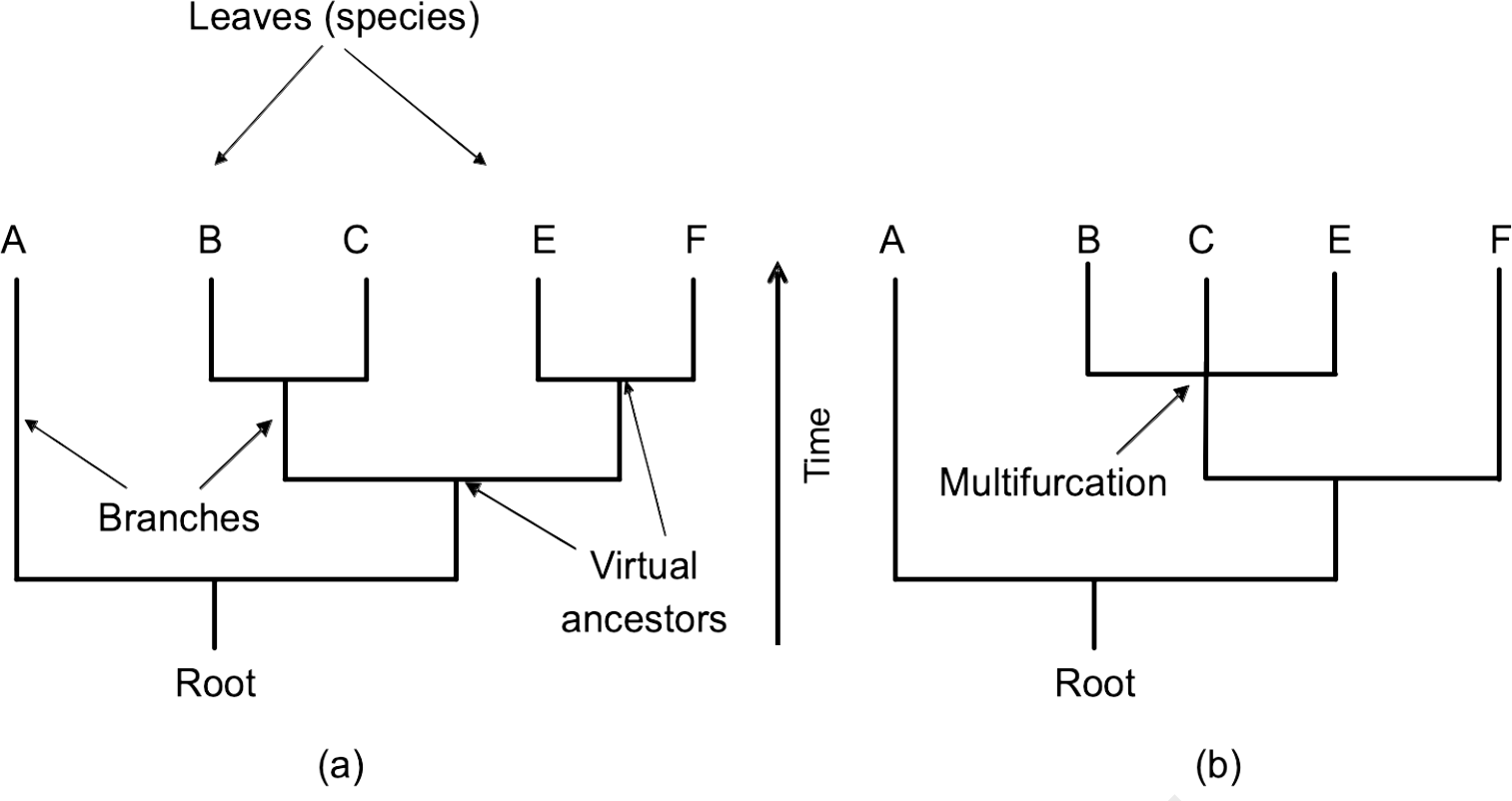
Illustration of phylogenetic tree with five species and resolved. The phylogenetic tree contains one root, virtual ancestors, branches and actual species (leaves).

### Robinson and Foulds topological distance

The Robinson and Foulds (RF) topological distance (4) between two trees is the minimum number of elementary operations (contraction and expansion) of nodes required to transform one phylogenetic tree into another. The two phylogenetic trees in question must have the same set of taxa. However, if the two trees do not contain the same set of taxa, a prepossessing step, consisting of the extraction of subtrees with common taxa, will be required. The more two phylogenetic trees are topologically close, the smaller is the value of the RF distance. However, the absolute value of RF does not take into account the number of taxa. It is often relevant to normalize this value by the maximum possible value of RF (equal to 2*n* − 6) for two binary trees with *n* leaves. This distance is mostly used in phylogenetic analysis.

### Consensus trees

Figure 2 presents an example of a set of four phylogenetic trees defined on a set of seven species and each tree is resolved. Many methods have been proposed to infer a single consensus tree for a collection of phylogenetic trees (6). The most known types of consensus trees are the strict consensus tree, the majority consensus tree and the extended majority consensus tree (2, 6). The strict consensus tree contains only the edges that are common to all input trees. The majority consensus tree contains the edges that are present in more than 50% of the input trees, although higher percentages may also be considered. According to the extended majority rule, the consensus tree includes all of the majority edges to which compatible residual edges are added gradually, starting with the most frequent ones. Extended majority consensus trees are the most frequently used consensus trees in evolutionary biology because they are usually much better resolved (i.e., have the lower total degree of internal nodes) than strict and majority consensus trees (2).

1

**Fig. 2.**
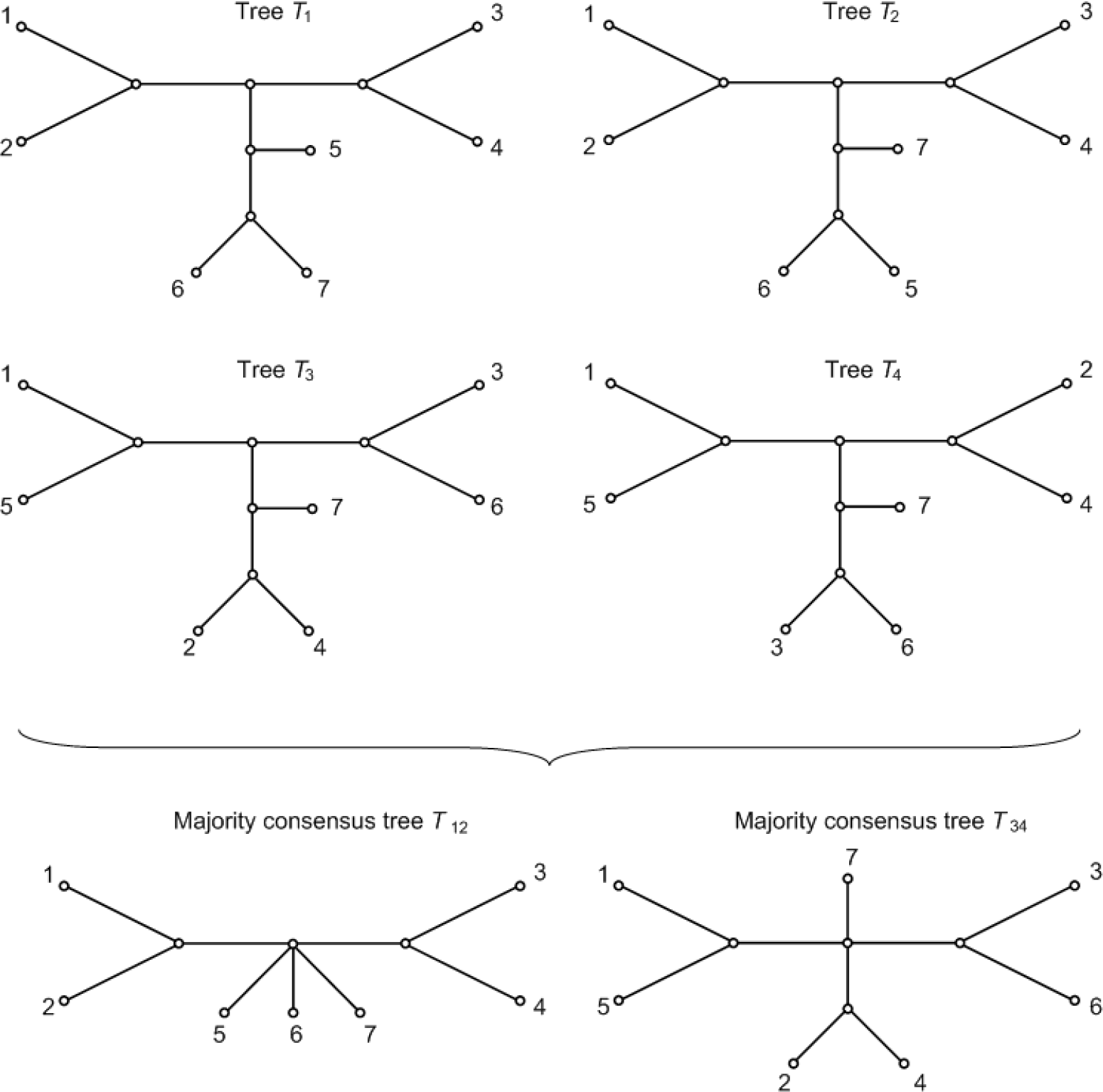
Four input phylogenetic trees defined with seven same species.

We can observe on Figure 2 that trees *T*_1_ and *T*_2_ are mostly similar and trees *T*_3_ and *T*_4_ are similar too. However, the extended majority consensus tree obtained by the set of input trees given by Figure 2 is represented by star tree. It would be more relevant to have not one consensus tree but two consensus trees.

## Data description

We will describe the sets of data simulated in our study, and we will present in detail the results of our simulations. We did three sets of simulations that are as follows:

### Simulation design

We tested our new algorithm for computing multiple consensus and supertrees. The simulation protocol include three main steps. First step, we randomly generated K consensus phylogenetic trees *T*_1_,; *T*_K_ with *n* leaves each, where K = 1, …, 5, and *n* = 8, 16, 32, 64 or 128. In the second step, for each phylogenetic tree *T_i_* (*i* = 1, …, K) obtained in the first step, we randomly generated a set of 100 trees 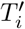 corresponding to cluster *i* and each 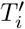 is differ from the Ti tree by two species switched. We classified each 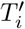 in four intervals of noise defined below. Each element 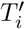 of *i* was a phylogenetic tree, such that the percentage of similarity (measured using the RF distance) between 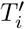 and *T_i_* varied from 0% to 10% (noise level 10% in Fig. 2c), from 10% to 25% (noise level 25%), from 25% to 50% (noise level 50%) or from 50% to 75% (noise level 75%). The third step consisted of a random removal of some species (the branches adjacent to these species were also removed) from the generated trees. The following percentages of species were removed: 0%, 10%, 25% or 50% (see Fig. 2). In the case of incomplete trees, the RF distance was computed between the maximum subtrees of two trees having at least 4 or more identical leaves. In the case of supertree, we filtered the two leaves set of two input trees for having the exactly all species in common. Thus, we also evaluated the ability of our algorithm to cope with incomplete data. The *k*-means algorithm was carried out with 100 random starts and until convergence of criterion score or maximum of 100 iterations in its internal loop. Note, for the case of consensus trees (see Fig. 2), we realized only the two first steps and add the third step for the case of supertrees. We illustrated only the results of CH criterion which is the best criterion when *k*=1 (see Fig. 3) and the criterion of supertree approach otherwise (see Fig. 4).

**Fig. 3.**
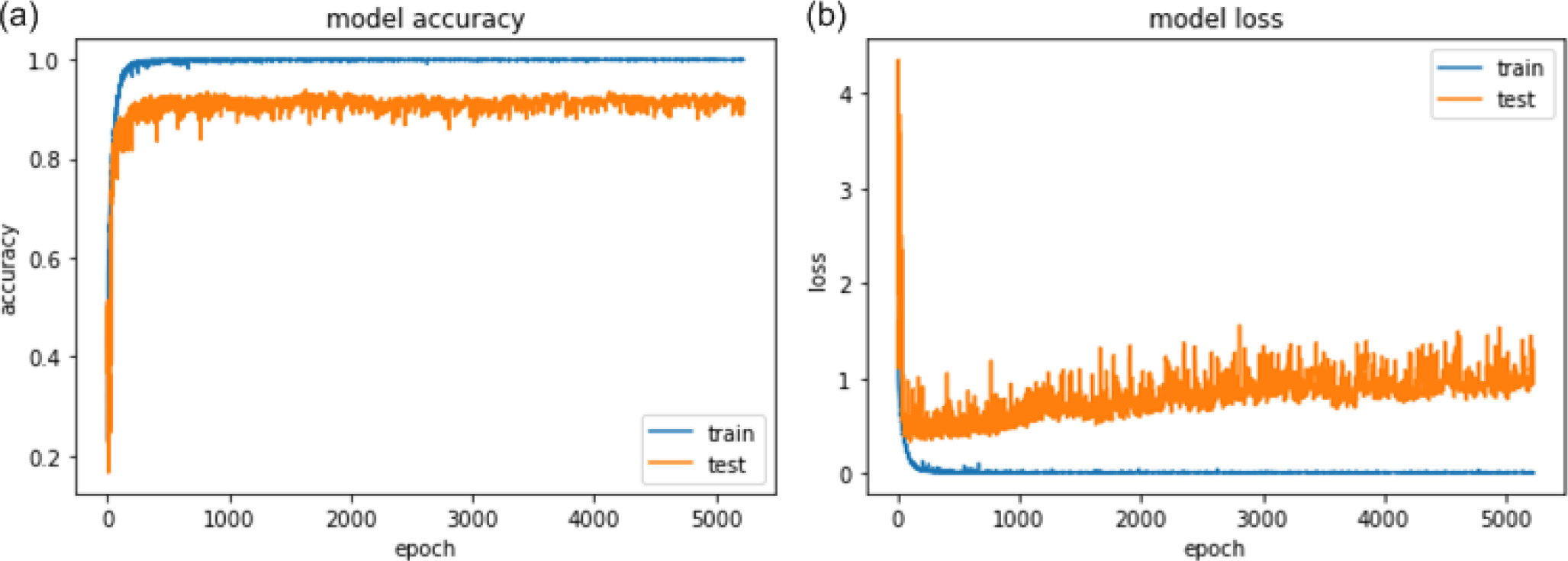
(a) A plot of model accuracy on train and validation datasets and (b) A plot of model loss on train and validation datasets.

**Fig. 4.**
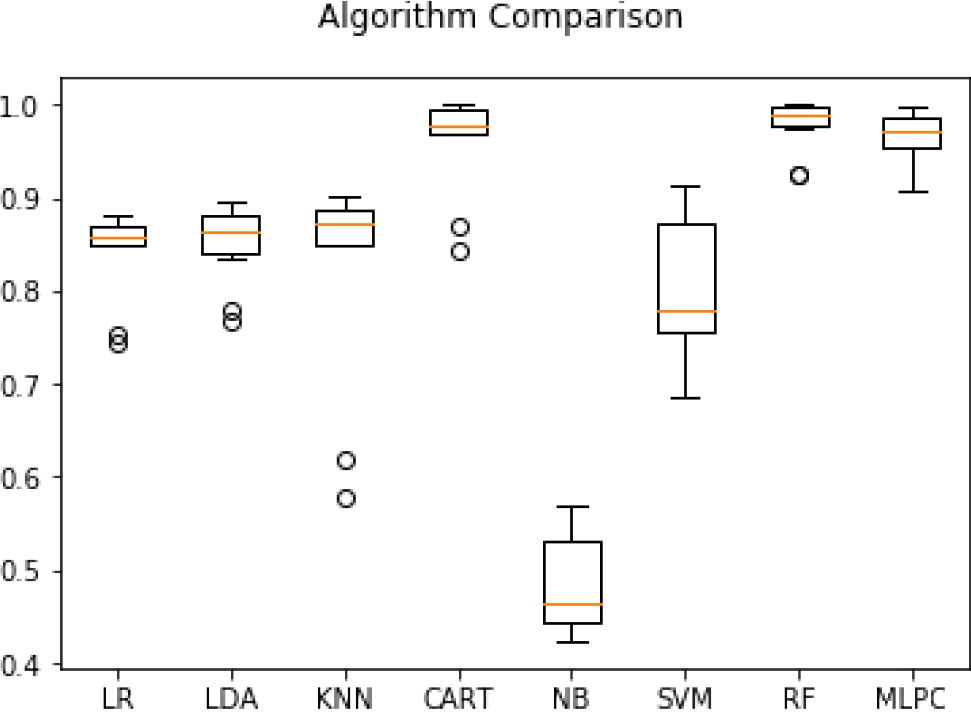
Box and Whisker Plots Comparing Algorithm Performance of eight models (1-Logistic Regression (LR), 2-Linear Discriminant Analysis (LDA), 3-K-Neighbors Classifier (KNN), 4-Decision Tree Classifier (CART), 5-GaussianNB (NB), 6-SVC (SVM), 7-Random Forest Classifier (RF), 8-MLP Classifier (MLPC)).

**Fig. 5.**
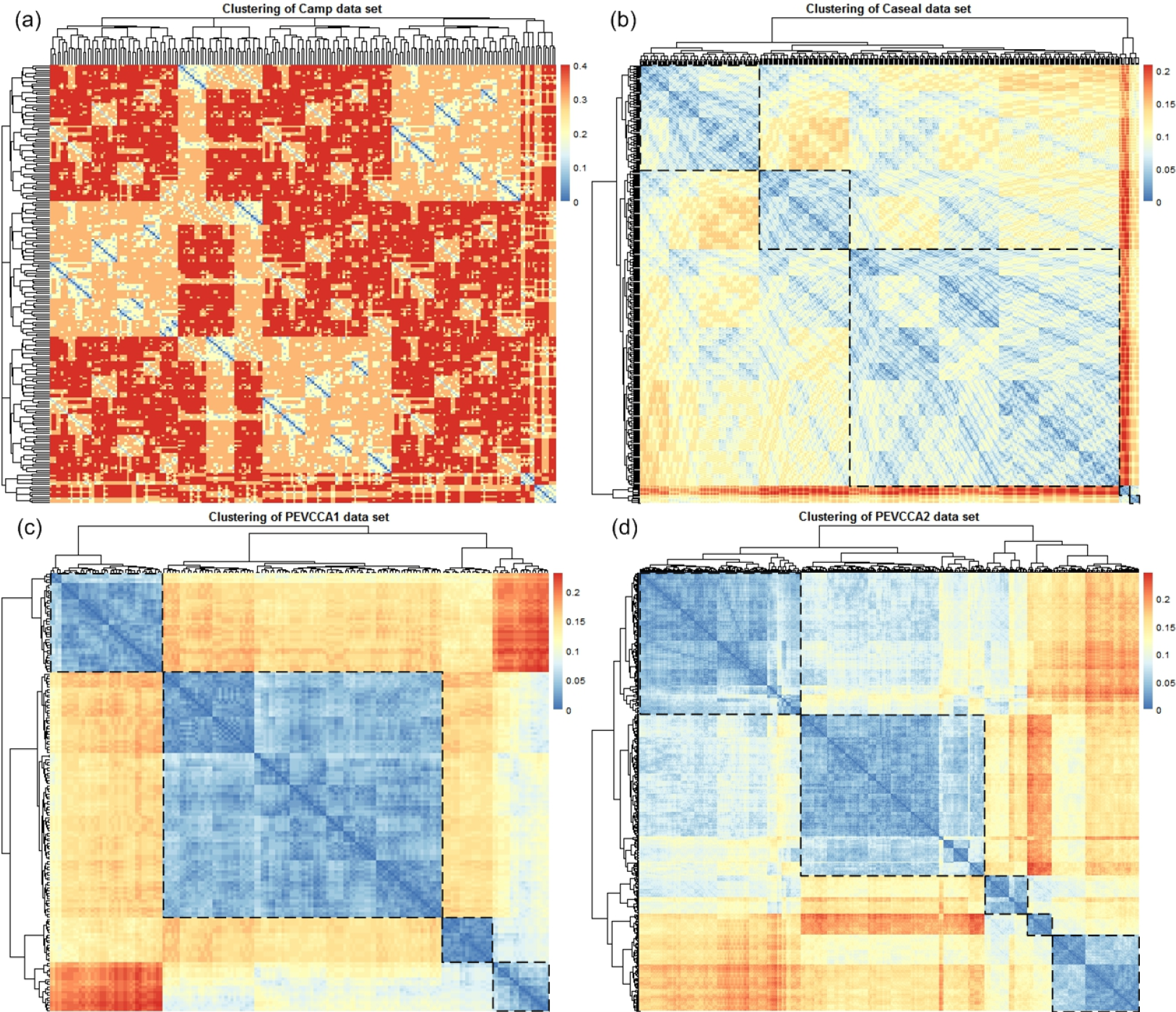
Heatmaps (a) of Camp dataset, (b) of Caesal dataset, (c) of PEVCCA1 dataset and (d) of PEVCCA2 dataset.

### Linguistic dataset

We extended our approach to linguistic dataset. The biologists are not alone to use the concept of evolutionary tree of Darwin by representing the histories of descent with modification. In fact, there are many studies which used the trees to study linguistic evolutionary by connecting the concepts of phylogeny and linguistic (Schleicher, 1873). More recently, this curious connection was presented by Atkinson and Gray (2005); they realized conceptual parallels between biological and linguistic evolution such as Horizontal gene transfer by borrowing and plant hybrids by language Creoles. We focused exclusively on the North and West Germanic groups. Among the 1315 word trees download and preprocessing from http://www.trex.uqam.ca/bioling_interactive/, data are also available from the github repository in https://github.com/TahiriNadia/CKMeansTreeClustering. Note, there are cases where Indo-European languages have more than one hypothesis of evolution. If this language belongs to the 12 languages studied (i.e North and West Germanic) then we have indicated on the tree of evolution the different hypotheses; otherwise we have masked these assumptions. We selected only the trees containing at least four languages of the North and West Germanic groups (i.e. Iceland-ic, Faroese, Swedish, Danish and Riksmål for North Germanic and Ducth, Flemich, Germanic ST, Frisian, PennDutch, Sranan and English for West Germanic), i.e. 248 trees in this step. We keep all the possibilities of evolutions concerning the languages of interest (i.e Sranan and Frisian). Then, we obtained 264 trees with 12 leaves.

### Stockham et al. datasets

To validate our approach, we used four biological datasets from (5) and compared our solutions to their results.

- The first dataset is Camp (i.e., Campanulaceae) (5, 7), which is a family of plants. This dataset is based on the deoxyribonucleic acid sequences of chloroplasts. These sequences are highly conserved (e.g., an order of genes, genome size), which is why they are frequently used as phylogenetic markers (8).
- The second set of dataset is Caesal (i.e., Caesalplinia) (5), which is included in the Angiosperms group. This dataset is based on the trnL-trnF intron and the chloroplast genome spacing regions.
- The third and fourth datasets are composed of PEVCCA1 and PEVCCA2 (i.e., Porifera, Echinodermata, Vertebrata, Cnidaria, Crustacea, and Annelida). These data are based on small ribosomal ribonucleic acid sequences of 129 species (5).

## Materials and Methods

In this section, we describe the main algorithm allowing us to classify a set of phylogenetic trees, defined on the same set of species using CNN approach.

In our simulations, we first generated random phylogenetic trees using the tree generation algorithm available on the TREX website (9). This algorithm requires as input the size of the tree, *n*, as well as the number of trees, *k*, and returns as output *K* random binary phylogenetic trees with *n* leaves each that is built according to the method of (10). In total, we generated 5808 unrooted phylogenetic trees for each of the following phylogenetic tree sizes: *n* = 8 and *n* = 16. Then, we computed the pairwise RF distances between trees. Finally, we normalized the matrix of RF distances by 2*n* − 6. After this last step, the values of RF will be between 0 and 1. We develop a novel DL architecture aimed at effectively including the phylogenetic structure into the learning process. The core of the network is CNN layers coupling with Keras library. The input layer is represented by a collection of a set of phylogenetic trees, specifically, the matrix of RF distance normalized. We activated input layer by sigmoid function. The neighbor detection procedure identifies the k-nearest neighbors of a given set of trees to be convolved with the filters by the CNN. To deal with the problem of finding neighbors for set of trees, we map the discrete space of the set of trees into an *L*_1_ norm (4). The selected neighbors are then convolved with the 20 filters on the CNN. We added seven hidden layers with Dropout at 0.1 (see Figure ??), we activated the neurons by Rectified Linear Unit (ReLU) function. The output layer are composed by MaxPooling, then Flatten layer, Fully Connected (Dense) for the transfer learning experiments and we used the same Dropout at 0.1. The neurons of the last layer were activated by softmax function. Finally, we used Adam to short for Adaptive Moment Estimation as optimizer with learning rate 0.01. The script was written in Python 3.6.

## Statistics

We present the obtained results using proposed method in this section. As well as the metrics (see Equations 1-4) that are utilized to evaluate the performance of methods.

### Statistic score

The *accuracy* of a test is its capability to recognize the classes properly. To evaluate the accuracy of the model, we should define the percentage of true positive and true negative in all estimated cases, i.e. the sum of true positive, true negative, false positive, and false negative. Statistically, this metric can be identified as follow:

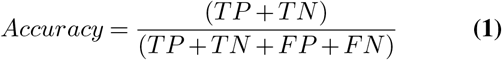

where *TP* is True Positive, *FP* is False Positive, *TN* is True Negative, and *FN* is False Negative.

The *precision* is a description of random errors, a measure of statistical variability. The formula of precision is the ratio between *TP* with all truth data (positive or negative).

The Equation is described as follow:

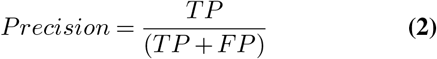

The *recall* or *sensitivity* or *TPRate* is defined as the number of true positive data labeled divided by the total number of *TP* and *FN* labeled data.

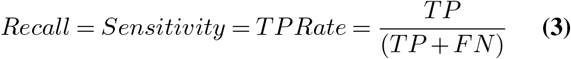

The *F* – *measure* or *F*_1_ is a well-known and reliable evaluation metric. The value of 1 would the mean perfect accuracy, i.e., the product would definitely be purchased.

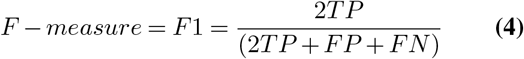

We examined these four evaluation metrics in our study (see the next section for the results of the *F*_1_ measure).

## Results

In this section, we will present the results of our simulations. These simulations will allow us to test the performance of our algorithm and evaluate the quality of our model. We consider two types of data: 1) The first type of data for which we have *a prior* knowledge of the number of clusters (i.e., simulated datasets), and 2) the second type of data for which the number of clusters is unknown initially, i.e., real datasets of (5) was considered (see Section “Data description”).

1

The results of accuracy and loss functions are shown in Figure 3. We performed 5227 epochs to reach the stability. Figure 3(a) shows the model is adequately chosen for your study. The accuracy of the train set is 0.9997 at 150 epochs and the accuracy of the test set is 0.8999 at 150 epochs.

Figure 3(b) shows the tendencies of loss function by epochs. The loss of the train set is 0.0047 at 150 epochs and the loss of the test set is 0.5195 at 150 epochs.

The obtained results demonstrate that the CNN approach to classifying the phylogenetic trees in the previous section clearly outperformed Stockham (5) approach.

### Simulated dataset

A list of each algorithm short name, the mean squared error and the standard deviation accuracy.

**Listing 1.**
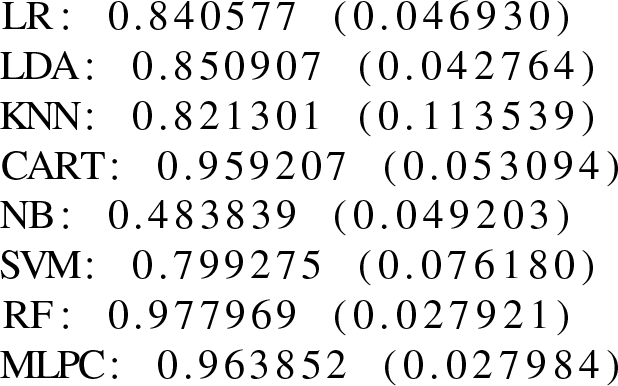
Output of comparing multiple algorithms.

Table 1 shows the best accuracy for a Random Forest function was the most accurate to the simulated dataset.

**Table 1.** Composition of North and West Germanic clusters and the results for supertree approach.

| Model Predictive | Accuracy |
| --- | --- |
| Logistic Regression | 85.37% |
| Support Vector Machine | 79.52% |
| Random Forest | 99.66% |
| Neural Network | 86.23% |
| Linear Discriminant Analysis | 86.06% |
| K-Neighbors Classifier | 83.82% |
| Decision Tree Classifier | 96.76% |
| Gaussian NB | 49.23% |

### Linguistic dataset

Our algorithm gives, for these input trees, three supertrees (See Table 2). The first cluster contained 76 trees, the second cluster contained 82 trees and the third cluster contained 106 trees.

For each of these three clusters, we used the software CLANN (Creevey and McInerney, 2004) and PAUP* (Swofford et al. 1993) to infer three supertrees (Fig. 8), one supertree by cluster obtained. This study shows that Riksmål language is the same evolution to Danish language (see Fig. 8b and 8c) and to Icelandic language (sees Fig. 8a). The same result has been obtained by Bryant et al. (2005), Gray et al. (2010) and Willems et al. (2016), showing that Riksmål lan-guage is a hybrid language between Icelandic and Danish languages. Riksmål is the most widely used written standard of contemporary Norwegian. Trees in Figure 8(a) identifies the branch of the West Scandinavian languages (Icelandic, Faroese and Norwegian), which constitutes one of the three branches of the North Germanic languages (Willems et al. 2016). However, Trees in Figures 8(b) and 8(c) highlights the influence of Danish on Riksmål. This influence is due to the political domination of Denmark over Norway between the end of the 14th century and the beginning of the 19th century. This study also shows that Sranan language is the same evolution to English language (see Fig. 8a and 8c) and to the clade (Dutch and Flemish) language (see Fig. 8b). The same result has been obtained by Bryant et al. (2005), Gray et al. (2010) and Willems et al. (2016), showing that Sranan language is hybrid language between English and Old Dutch languages. We also detect-ed that PennDutch language is hybrid language between clade of English and Sranan languages (see Fig. 8a and 8c) and German language (see Fig. 8b). Real evolutionary data often contain a number of different and sometimes conflicting phylogenetic signals, and thus do not always clearly support a unique tree. To address this problem, Bandelt and Dress (1992) developed the method of split decomposition. The split decomposition method introduced by decomposes the given distance matrix into simple component based on weighted splits Bandelt and Dress (1992). For summarise a set of trees, we used SplitsTree4 (Huson and Bryant 2005) with all North and West Germanic dataset after preprocessing (see Fig. 7a), with the three supertrees obtained by our new algorithm wit supertree approach (see Fig. 7b) and with each cluster obtained by supertree approach (see Fig.9 in Appendices).

We extended our approach to linguistic dataset. The biologists are not alone to use the concept of evolutionary tree of Darwin by representing the histories of descent with modification. In fact, there are many studies which used the trees to study linguistic evolutionary by connecting the concepts of phylogeny and linguistic (Schleicher, 1873). More recently, this curious connection was presented by Atkinson and Gray (2005); they realized conceptual parallels between biological and linguistic evolution such as Horizontal gene transfer by borrowing and plant hybrids by language Creoles. We focused exclusively on the North and West Germanic groups. Among the 1315 word trees download and preprocessing from http://www.trex.uqam.ca/bioling_interactive/, data are also available from the github repository in https://github.com/TahiriNadia/CKMeansTreeClustering. Note, there are cases where Indo-European languages have more than one hypothesis of evolution. If this language belongs to the 12 languages studied (i.e North and West Germanic) then we have indicated on the tree of evolution the different hypotheses; otherwise we have masked these assumptions. We selected only the trees containing at least four languages of the North and West Germanic groups (i.e. Icelandic, Faroese, Swedish, Danish and Riksmål for North Germanic and Ducth, Flemich, Germanic ST, Frisian, PennDutch, Sranan and English for West Germanic), i.e. 248 trees in this step. We keep all the possibilities of evolutions concerning the languages of interest (i.e Sranan and Frisian). Then, we obtained 264 trees with 12 leaves.

**Stockham et al. datasets. 1**

## Discussion

In this article we described a new algorithm for partitioning a set of phylogenetic trees in several clusters in order to infer multiple supertrees, in case, the input trees have different, but mutually overlapping sets of leaves. We presented new formulas allowing for using the popular Calin’ski-Harabasz, Silhouette and Gap statistic cluster validity indices as well as the Robinson and Foulds topological distance in the framework of tree clustering based on the popular k-means algorithm. The new algorithm can be used to address a number of important issues in bioinfor-matics, such as the identification of genes having similar evolutionary histories, e.g. those that underwent the same horizontal gene transfers or those that were affected by the same ancient duplication events. It can also be used for inference of multiple subtrees of Tree of Life. In order to compute the Robinson and Foulds topological distance between such pairs of trees, we can first reduce them to the common set of leaves. After this reduction, the Robinson and Foulds distance normalized by its maximum value, which is equal to 2*n*-6 for two binary trees with *n* leaves. Overall, good performances achieved by the new algorithm in terms of both clustering quality and running time makes it well suited for the analysis of large genomic and phylogenetic datasets. A Python script, called KMSTC (K-Means SuperTree Clustering), implementing the discussed tree partitioning algorithm is freely available at https://github.com/TahiriNadia/KMeansSuperTreeClustering/.

## Supporting Information

If you intend to keep supporting files separately you can do so and just provide figure c aptions h ere. Optionally make clicky links to the online file using `\href{url}{description}`.

## Availability of data and materials

All the data presented in this article and the code are available at: https://github.com/TahiriNadia/ML_DL_Classification_Trees.

## Acknowledgments

We thank B. Mazoure of Montreal Institute for Learning Algorithms (MILA) for critical reading of the article.

## Supplementary Note 1: ANNEXE

**Table 2.**
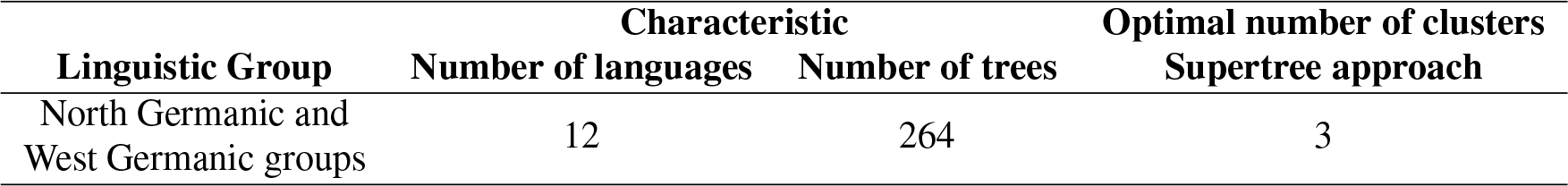
Composition of North and West Germanic clusters and the results for supertree approach.

| Linguistic Group | Characteristic |  | Optimal number of clusters<br>Supertree approach |
| --- | --- | --- | --- |
|  | Number of languages | Number of trees |  |
| North Germanic and<br>West Germanic groups | 12 | 264 | 3 |

